# T Cell Receptor Non-Equilibrium Kinetics

**DOI:** 10.1101/2021.10.27.466112

**Authors:** Zachary A. Rollins, R. Faller, Steven C. George

## Abstract

An atomic-scale mechanism describing the role of mechanosensing in T Cell Receptor (TCR) recognition of peptides in the binding groove of the peptide-major histocompatibility complex (pMHC) may inform the design of novel TCRs for immunotherapies. Using steered molecular dynamic simulations, our study demonstrates that mutations to peptides in the binding groove of the pMHC – which are known to discretely alter the T cell response to an antigen – influence MHC conformation and thus the overall strength of the TCR-pMHC bond including duration and length under constant load. Moreover, physiochemical features of the TCR-pMHC dynamic bond strength, such as hydrogen bonds and Lennard-Jones contacts, correlate with the immunogenic response elicited by the specific peptide in the MHC groove. Thus, formation of transient TCR-pMHC bonds is a characteristic of immunogenic peptides and is mediated by stabilized interactions.x

## INTRODUCTION

T cells discriminate peptide-major histocompatibility complexes (pMHCs) expressed on antigen presenting cells (APCs) via their T Cell Receptors (TCRs). Engineered T cells have elicited exciting clinical responses in cancer patients(1–5); however, off-target toxicity is a prevailing limitation and has created a need for a deeper understanding of how the TCR engages its antigen and induces an immunogenic response.

Several features of the TCR-pMHC interaction have been evaluated in an effort to identify quantitative descriptors that predict *in vitro* T cell immunogenicity(6–8). For example, equilibrium TCR-pMHC kinetic parameters, measured by surface plasmon resonance, have generated a collection of theories to explain specificity including kinetic proofreading(9–12), serial triggering(13–15), and the law of mass action(16–19). However, none of these theories can completely predict the immunogenic response across multiple TCR-pMHC systems(20–22), indicating that equilibrium-based models are insufficient.

*In vivo* binding of a cytotoxic T cell to an APC is a dynamic process that introduces mechanical forces on the TCR-pMHC interaction(23, 24). Using DNA-based tension probes, mechanical forces on the TCR-pMHC bond have been estimated in the 10-20 pN range(25, 26). These tensile forces may be attributed to interstitial shear on cells(27), relative membrane sliding during cell motility(28), or actin polymerization during formation of the immunological synapse(29, 30). To investigate the effects of these forces, several biophysical techniques—including the biomembrane force probe(31–35), optical tweezers(36), and the laminar flow chamber(37, 38) —have been used to characterize the force-dependent TCR-pMHC dissociation kinetics. Initial reports demonstrate that bond lifetimes increase with applied force (“catch”) until a critical force (~10-15 pN) is reached, and then decrease with further applied force (“slip”). This “catch-slip” bond behavior has only been reported for TCR-pMHCs that generate an immunogenic response (e.g. *in vitro* INFγ production)(31–35). Consequently, this catch-slip bond behavior has been postulated as a mechanotransduction discrimination mechanism determining T cell immunogenicity.(39) However, a more recent report(38) using a laminar flow chamber found no catch-slip bond behavior even for immunogenic TCRs(40), but did demonstrate a positive correlation between bond strength and immunogenicity. These studies clearly demonstrate that dissociation kinetics at critical forces (~10-20 pN) correlate with immunogenicity(31–38), highlighting the importance of TCR-pMHC bond strength, and suggest a force-dependent kinetic proofreading discrimination model. In such a model, the TCR would sustain and form transient bonds under load for sufficient time to initiate biochemical signaling. While intriguing, the potential underlying molecular level mechanisms for bond lifetime enhancement under load are unknown.

Emergence of mechanosensing by the T cell underscores the importance of the dynamic energy landscape and molecular mechanics of the TCR-pMHC interaction under tensile force. Steered molecular dynamics (SMD) simulations have been used to investigate these properties(34, 35, 41). These reports have argued that transiently formed Lennard-Jones contacts (LJ-Contacts) or hydrogen bonds (H-Bonds) are responsible for increased bond lifetime. In contrast, a recent study argues that the constant regions of the TCR stabilize and preserve the interfacial pMHC H-Bonds and LJ-Contacts under physiological load (~10-20 pN). A comprehensive analysis that addresses these features is crucial to decipher the relative energetic contributions of transient, stabilized H-Bonds and LJ-Contacts to the overall TCR-pMHC bond strength. Additionally, investigation into localized interactions as well as comparison of global and local essential motion will reveal more precise information on the structural and energetic mechanisms of TCR-pMHC interactions. Finally, atomic fluctuations in receptor-ligand interactions result in a stochastic ensemble of equilibrium configurations and the consequential heterogeneity of SMD simulation results remains unaddressed(42).

Here, we performed SMD in triplicate by applying a linear potential (i.e., a constant force) to the center of mass (COM) of the complex formed by the DMF5 TCR and pMHC bound to the peptide MART1(AAGIGILTV) (PDB ID: 3QDJ). We investigated the complex when pMHC is bound to the L1 (LAGIGILTV) and GVA (GAGIGVLTA, β-endoxylanase) peptide mutants, which have been reported as a super-agonist (increased immunogenic response compared to wildtype) (43) and antagonist (no immunogenic response)(44), respectively (**Figure 1, Supplementary Videos**). We found that the peptide identity alters the conformation of the binding groove in the MHC, and these conformational changes result in characteristic sets of H-Bonds and LJ-Contacts that significantly contribute to the overall bond strength. These sets of H-Bonds and LJ-Contacts are uniquely distributed across the pMHC interface, occur at characteristic time and distance scales, and correlate with the immunogenicity of the specific peptide. Furthermore, under load, all TCR-pMHCs maintain their secondary structure and essential atomic motion is predominantly in the direction of applied force. These results provide new atomic-level insight on TCR-pMHC bond strength.

**Figure 1:**
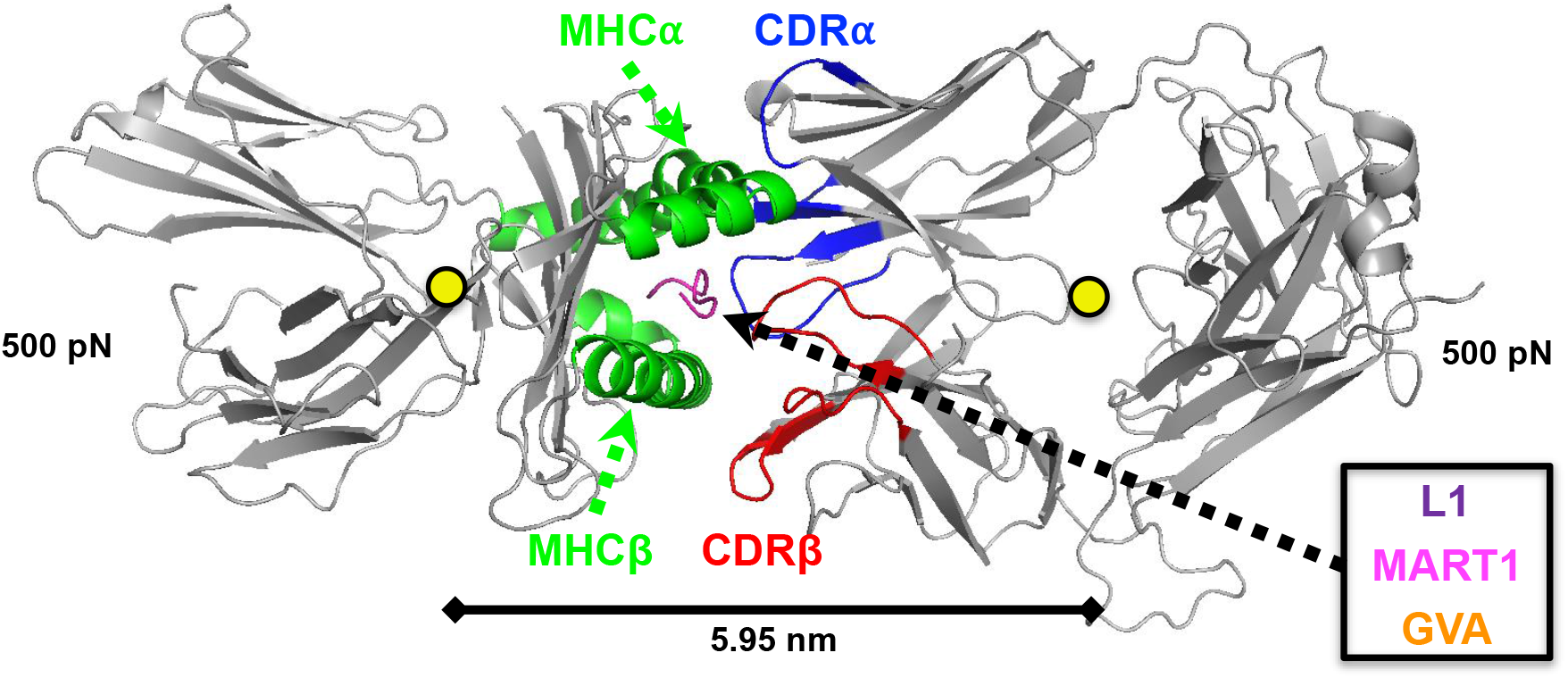
Graphic representation of Steered Molecular Dynamics (SMD) simulations on a TCR-pMHC system. This includes highlighted interfacial substructures [MHCa & MHCb = green, TCR CDRa = blue, TCR CDRb = red, Peptide (L1, MART1, GVA) = purple, magenta, orange], direction of pulling (yellow circles and arrows, respectively), and position of and distance between COMs (scale bar at bottom). The non-interacting bodies of the TCR and pMHC are colored in gray.

## METHODS

### Molecular Dynamics Setup

The crystal structure of the human DMF5 TCR complexed with agonist peptide MART1-HLA-A2 (PDB code: 3QDJ) was the initial structure for all simulations (45). Amino acid substitutions were made to the MART1 peptide (AAGIGILTV) using the Mutagenesis plugin on Pymol Molecular Graphics System (Schrodinger, LLC; New York, NY) to generate starting configurations for the L1 (**L**AGIGILTV) and GVA (**G**AGIG**V**LT**A**) peptides. Interfacial substructures were defined by sequential residues from the corresponding chains: TCRα (CDR1α: 24-32, CDR2α: 50-55, CDR3α: 89-99), TCRβ (CDR1β: 25-31, CDR2β: 51-58, CDR3β: 92-103), MHCα (MHCβ: 50-85, MHCα: 138-179), and peptide (L1: 1-9, MART1: 1-9, GVA:1-9). To determine protonation states, pKa values were calculated using propka3.1(46, 47) and residues were considered deprotonated in Gromacs(48) if pKa values were below physiologic pH 7.4. The resulting systems were solvated in rectangular water boxes using the TIP3P water model(49) large enough to satisfy the minimum image convention in order to avoid artifacts by self-interaction. Na^+^ and Cl^-^ ions were added to neutralize protein charge and reach physiologic salt concentration ~150 mM. All simulations were performed with Gromacs 2019.1(48) using the CHARM22 plus CMAP force field for proteins(50) and orthorhombic periodic boundary conditions. All simulations were in full atomistic detail.

### Energy Minimization and Equilibration

Generating equilibrated starting structures for the steered molecular dynamics simulations required four steps. (1) Steepest descent energy minimization to ensure correct geometry and the absence of steric clashes. (2) 100 ps simulation in the constant volume (NVT) ensemble to bring atoms to correct kinetic energies. A temperature of 310 K was maintained by coupling all protein and nonprotein atoms in separate baths using the velocity rescaled thermostat with a 0.1 ps time constant(51). (3) 100 ps simulation in the constant pressure (NPT) ensemble using Berendsen pressure coupling(52) and a 2.0 ps time constant to maintain isotropic pressure at 1.0 bar. (4) Production MD simulations were conducted for 100 ns with no restraints. Steps (2)-(3) used position restraints on all protein atoms. To ensure true NPT ensemble sampling during 100 ns production runs, the Nose-Hoover thermostat(53) and Parrinello-Rahman barostat(54) were used to maintain temperature and pressure with time constants2.0 and 1.0ps, respectively, utilizing the isothermal compressibility of water, 4.5^-5^ bar^-1^. Box size for equilibration was 10.627 x 7.973 x.10.685 nm^3^ with ~ 48,000 water molecules, ~300 ions, and ~157,000 total atoms (**Table 1**). All simulations used the Particle Ewald Mesh algorithm(55, 56) for long-range electrostatic calculations with cubic interpolation and 0.12 nm maximum grid spacing. Short-range nonbonded interactions were cut off at 1.2 nm. All water bond lengths were constrained with SETTLE(57) and all other bond lengths were constrained using the LINCS algorithm(58). The leap-frog algorithm was used for integrating equations of motion with a 2 fs time steps. After 100, 95, and 90 ns of production run, respectively, MD configurations for each peptide mutant were extracted and used as the three initial configurations for steered molecular dynamics simulations.

**Table 1:** Equilibration of mutant peptides including amino acid sequence, number of solvent molecules, number of ions, and total atoms in respective simulation boxes

| TCR-pMHC | Peptide Amino Acid Sequence | Equilibration Water Molecules | Equilibration Ions | Equilibration Total Atoms |
| --- | --- | --- | --- | --- |
| L1 | LAGIGILTV | 47998 | 164 Na, 144 Cl | 157134 |
| MART1 | AAGIGILTV | 48001 | 164 Na, 144 Cl | 157134 |
| GVA | GAGIGVLTA | 47997 | 164 Na, 144 Cl | 157110 |

### Steered Molecular Dynamics (SMD)

The full protein structure was extracted for each peptide mutant (L1, MART1, GVA) at the above-described production MD configurations (100, 95, 90 ns). These protein complexes were aligned along the *x*-axis and solvated in rectangular TIP3P water boxes with dimensions 30 x 9.972 x 12.685 nm^3^ and Na ^+^ and Cl^-^ ions were added to neutralize protein charge and reach ~150 mM sat concentration (see **Table 2**). Before pulling, all systems underwent (1) energy minimization (2) 100 ps NVT (3) and 100 ps NPT. During pull, the Nose-Hoover thermostat and Parrinello-Rahman barostat were again used to maintain temperature and pressure. A 500 pN linear potential was applied to the center of mass (COM) of the TCR and pMHC in the *x*-direction and simulations continued until the distance between COMs reached 0.49 times the box size. The COM was chosen as the site of applied force because pulling from the TCR and MHC termini results in artificial unfolding (**Figure S1**). The termini control simulation was performed on the 100 ns MART1 configuration, with equivalent force magnitude (i.e., 500 pN), and was pulled from the COM of the terminal chain residues (MHCα: 275, TCRα: 199 & TCRβ: 242) (**Figure S1**). Furthermore, while observing bond lifetimes at a force of 500 pN is computationally feasible, we understand that this is higher than the physiological force range. Thus, we demonstrated, for a series of configurations along the reaction coordinate, that the root mean square fluctuations of the TCR-pMHC interface are not different at a physiological separation force of 10 pN (**Figure S2**). These force control simulations (i.e., 10 pN) were pulled from the COM and performed for 50 ns on multiple configurations along the reaction coordinate (i.e., 6.16 and 6.20 nm) of the 100 ns MART1 configuration 500 pN pull. All simulation trajectories and selected frames visualized using the Pymol Molecular Graphics System (Schrodinger, LLC; New York, NY).

**Table 2.** Steered MD simulations of mutant peptides including amino acid sequence, number of solvent molecules, number of ions, and total atoms in respective simulation boxes

| TCR-pMHC | Peptide Amino Acid Sequence | Equilibration Time (ns) | SMD Water Molecules | SMD Ions | SMD Total Atoms |
| --- | --- | --- | --- | --- | --- |
| L1 | **LAGIGILTV** | 100 | 118977 | 363 Na,<br>343 Cl | 370469 |
|  |  | 95 | 118993 | 363 Na,<br>343 Cl | 370517 |
|  |  | 90 | 118960 | 363 Na,<br>343 Cl | 370418 |
| MART1 | **AAGIGILTV** | 100 | 118968 | 363 Na,<br>343 Cl | 370433 |
|  |  | 95 | 118925 | 363 Na,<br>343 Cl | 370304 |
|  |  | 90 | 118934 | 363 Na,<br>343 Cl | 370331 |
| GVA | **GAGIGVLTA** | 100 | 118977 | 363 Na,<br>343 Cl | 370448 |
|  |  | 95 | 119013 | 363 Na,<br>343 Cl | 370556 |
|  |  | 90 | 118973 | 363 Na,<br>343 Cl | 370436 |

### Endpoints and Data Analysis

Data analysis was performed by built in Gromacs functions, standard python packages for data handling and visualization (i.e. numpy, pandas, seaborn, matplotlib, statistics, and GromacsWrapper), and custom python scripts. The geometry of a Lennard-Jones contact is defined as a distance less than 3.5 nm between atoms. The principal component analysis of the simulation trajectories was performed using the package MDAnalysis(59, 60). Secondary structure was designated based on a hydrogen bond estimation algorithm(61, 62). Custom scripts relevant to the production of figures have been made available on a Github repository: https://github.com/zrollins/TCR-pMHC.git.

### Statistical Analysis

All data was processed in python utilizing standard packages for data handling and visualization (i.e. numpy, pandas, seaborn, matplotlib, statistics, scipy, and pingouin). Results were presented as mean ± SEM. As indicated in figures, statistics were performed in python using scipy for Student’s t-tests, scipy for one-way analysis of variance (ANOVA), and pingouin for pairwise Tukey-HSD post-hoc tests. Detailed outputs of statistical analysis were written to excel and are provided as supporting information.

## RESULTS & DISCUSSION

### Equilibrated Structures and Potential Mean Force

We initiated our simulation with the crystal structure of the TCR-pMHC complex bound to the MART1 peptide (PDB ID: 3QDJ)(45); biomolecular simulations of systems this size require tens of nanoseconds to reach equilibration(63). TCR-pMHC complexes bound to the L1, MART1, and GVA peptides were simulated for 100 ns and reached equilibrated configurations around 60, 20, and 40 ns, respectively (**Figure S3 A-C**). After convergence, atomic fluctuations generate an ensemble of configurations demonstrated by the aligned equilibrated structures at 100, 95, and 90 ns for the respective mutants (**Figure 2A-C**). In the 90 to 100 ns time window, the root mean square fluctuations range from 0.05 - 0.10 nm for all TCR-pMHC interfacial substructures (**Figure 2D**). Although there are statistically significant differences in TCR substructure fluctuations (i.e., CDR1α, CDR2α, CDR3α, and CDR3β), there is no trend and 0.2 Å differences in fluctuation are unlikely to have physical significance.

**Figure 2:**
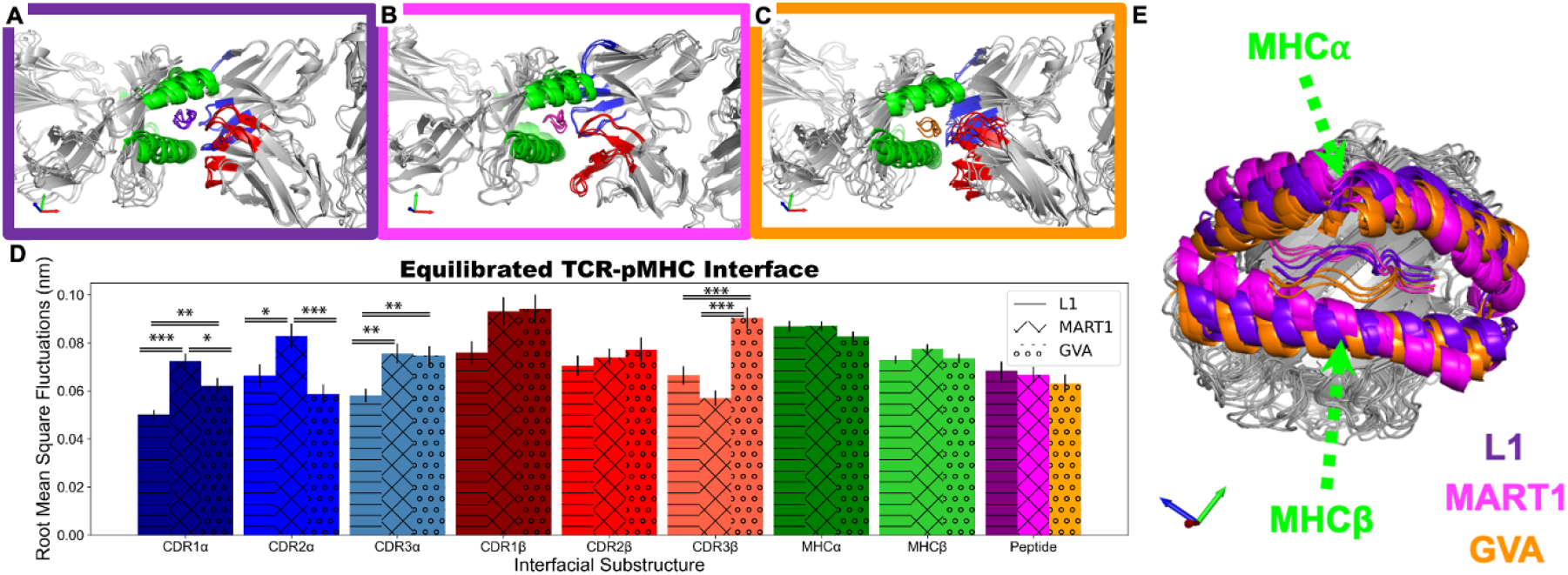
Equilibrated TCR-pMHC Structures from 100, 95, and 90 ns included with Interfacial Substructures. (**A**) L1 Peptide (purple) and TCR-pMHC (**B**) MART1 Peptide (magenta) and TCR-pMHC (**C**) GVA Peptide (orange) and TCR-pMHC (**D**) Interfacial substructure root mean square fluctuations for each TCR-pMHC from 90 to 100 ns (**E**) Aligned pMHC structure at 100, 95, and 90 ns for each peptide (L1=purple, MART1=magenta, GVA=orange). The equilibrated MHCa & MHCb are indicated in the peptide color to distinguish structure (previously green in A-C). Subsctructures of mutants were statisically compared (n=3): #p<0.10, *p<0.05, **p<0.01, ***p<0.001 by one-way ANOVA followed by Tukey-HSD post-hoc test.

Interestingly, mutations to the peptide determine the conformation of the surrounding MHC α-helices (**Figure 2E)** and TCR substructures (**Figure S4**) and are independent of fluctuations. These configurations were separated by applying a 500 pN linear potential to the COM of the TCR and pMHC for each of the three starting configurations determined at 100, 95, and 90 ns equilibration time. To broadly assess TCR-pMHC bond strength, we evaluated interaction energy, H-Bonds, and LJ-Contacts on COM distance and time scales. Error bars represent the standard error of measurement (SEM) for each peptide (L1, MART1, and GVA) over the three SMD simulations. In accordance with immunogenicity, longer reaction distance (**Figure 3A**) and non-linear energetic decay (**Figure 3A-B**) for L1 and MART1 indicate higher bond strength than GVA. Remarkably, structural alterations to the MHC α-helices (**Figure 2E**) determine TCR-MHC reaction distance (**Figure 3A**) for L1, MART1, and GVA. Therefore, the unique sequence of the peptide functions as the key that unlocks the TCR-pMHC interaction. This concurs with experimental observations of peptide altered MHC binding grooves(64) and peptide altered immunogenic response(43, 44). For receptor-ligand interactions, Coulombic and Lennard-Jones potential are dominated by H-Bonds and hydrophobic LJ-Contacts, respectively.

**Figure 3:**
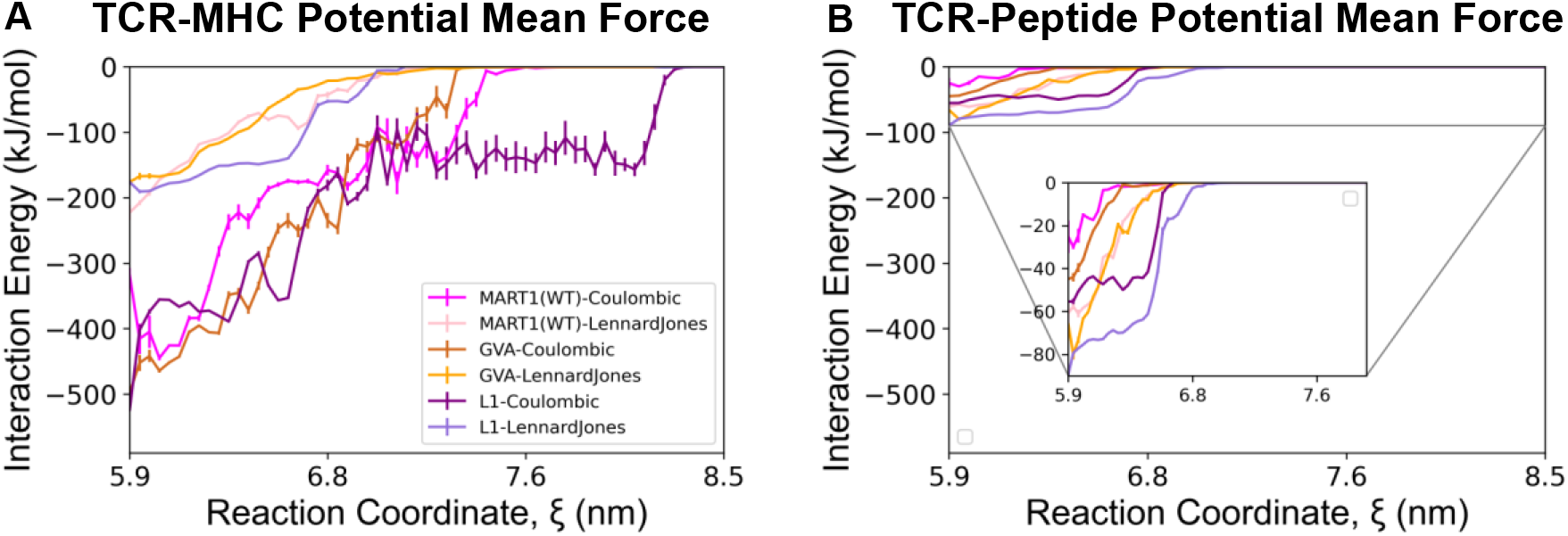
Potential Mean Force. (**A**) TCR-MHC and (**B**) TCR-Peptide interaction energy is separated into Lennard-Jones and Coulombic potential for each peptide and plotted against the reaction coordinate defined as the TCR-pMHC COM distance. For TCR-pMHC SMD simulations (100, 95, 90 ns), interaction energies were placed into ~0.5 Å bins and error represents SEM.

The TCR-MHC Coulombic potential is higher in magnitude than the Lennard-Jones potential for all mutants along the reaction coordinate (**Figure 3A).** Conversely, the TCR-Peptide Lennard-Jones potential is stronger than the Coulombic potential for all mutants along the reaction coordinate (**Figure 3B**). Thus, these results indicate that there is localized energetic heterogeneity, where H-Bonds are more consequential for TCR-MHC interactions and LJ-Contacts are more consequential for TCR-Peptide interactions. The general strength of the TCR-MHC interaction is substantially (5x) larger than the TCR-Peptide interaction.

### Hydrogen Bonds and Lennard-Jones Contacts

In order to gauge the importance of transient and stable H-Bonds or LJ-Contacts, we utilized probability densities and existence occupancies. Probability densities are defined for H-Bonds and peptide solvent accessible surface area (SASA). Existence occupancy is defined as the percentage of time under load that unique H-Bonds or LJ-Contacts exist; thus, higher occupancy cutoffs represent more stable interactions.

Higher expected value of H-Bonds from the probability density distribution (**Figure 4A**) for L1 and MART1 (5.99 and 5.04 compared to GVA 4.47) combined with more low occupancy H-Bonds for MART1 (**Figure 4B**) demonstrate the importance of transiently formed H-Bonds in TCR-pMHC bond strength. Additionally, the trend continues for more high occupancy (up to 10%) H-Bonds for L1 and MART1 demonstrating the importance of more stabilized interactions (**Figure 4B**); however, there is no statistical significance. No statistical significance of high occupancy H-Bonds between mutants is likely the result of one heterogenous sample for L1 and MART1 in the stochastic ensemble (n=3) demonstrating the importance of evaluating the heterogeneity of SMD results by performing several independent simulations. Furthermore, there is qualitative agreement between the Coulombic potential reaction distance (**Figure 3A-B**) and the number of hydrogen bonds for TCR-Peptide (**Figure S5Aii**) and TCR-MHC (**Figure S5Aiii**) interactions. In previous work with a different system, we have demonstrated that increased hydrophobic SASA yields greater aggregation propensity(65) and likely more hydrophobic interactions. Higher mean probability density of peptide hydrophobic SASA for L1 and MART1 suggest more hydrophobic interactions (**Figure 4C**). More numerous low (>0%) and high occupancy (up to 25%) LJ-Contacts for L1 and MART1 highlight the importance of transient and stabilized hydrophobic interactions (**Figure 4D**). Moreover, there is qualitative agreement between the Lennard Jones potential reaction distance (**Figure 3A-B**) and the number of LJ-Contacts for TCR-Peptide (**Figure S5Bii)** and TCR-MHC (**Figure S5Biii**) interactions. Probability densities and existence occupancies are more numerous under load for biologically immunogenic peptides. This evidence illustrates that bond strength is a consequence of stabilized and transiently formed H-Bonds and LJ-Contacts.

**Figure 4:**
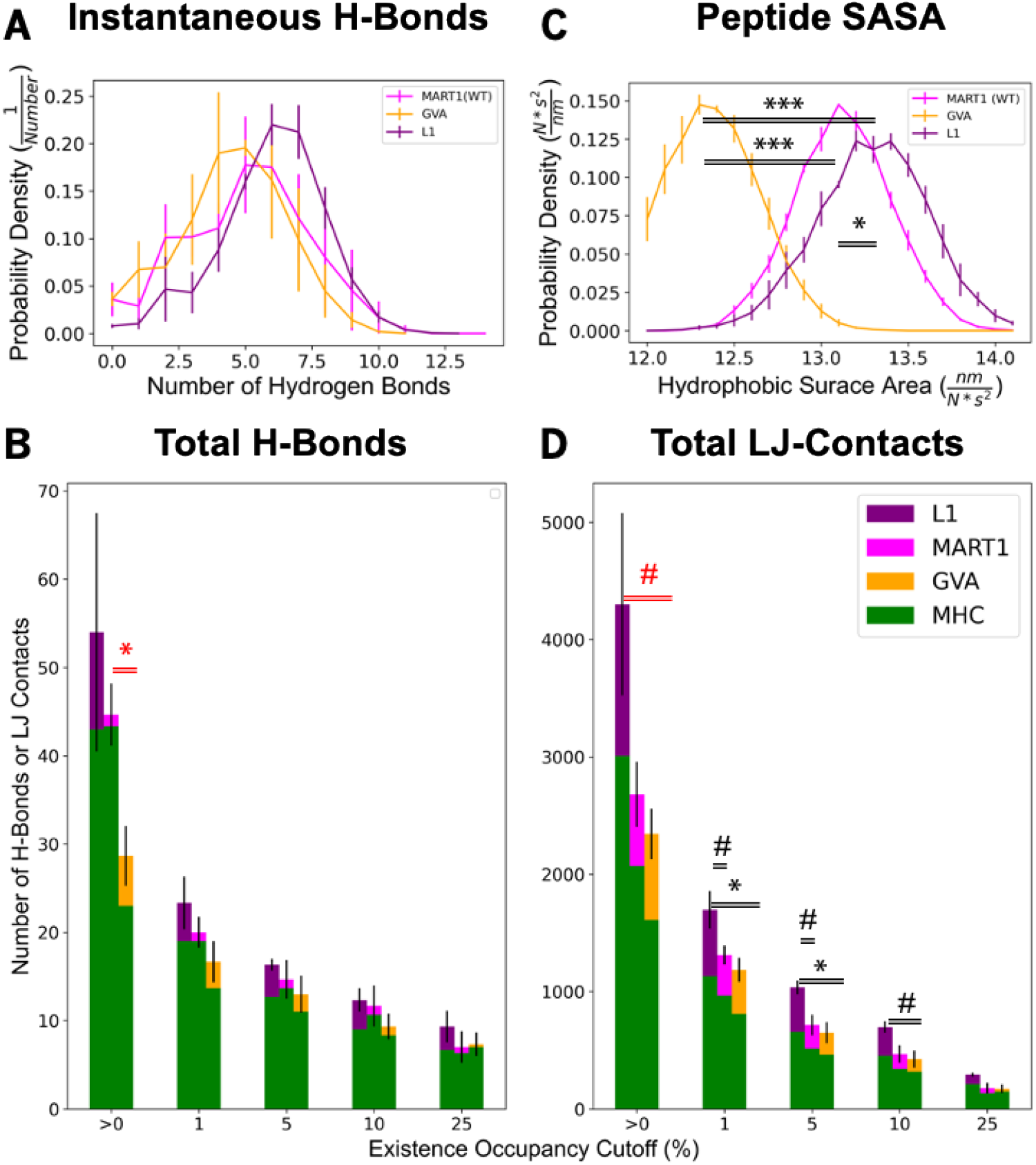
Hydrogen Bonds and Lennard-Jones Contacts. (**A**) For each peptide, probability density of hydrogen bonds between TCR and pMHC is plotted. For TCR-pMHC SMD simulations (100, 95, 90 ns), times (with 1 ps resolution) were distributed into number of hydrogen bond bins and error represents SEM. (**B**) Number of unique hydrogen bonds are graphed with increasing existence occupancy between the TCR and pMHC. The interactions are separated into TCR-MHC H-Bonds (green) and TCR-Peptide H-Bonds (L1=purple, MART1=magenta, GVA=orange) and stacked. Error represents SEM over three SMD simulations. **(C)** For each peptide, probability density of solvent accessible surface area (SASA) is plotted. For TCR-pMHC SMD simulations (100, 95, 90 ns), times (with 1 ps resolution) were cut into 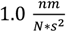 bins and error represents SEM. **(D)** Number of unique LJ-Contacts are graphed with increasing existence occupancy between the TCR and pMHC. The interactions are separated into TCR-MHC LJ-Contacts (green) and TCR-Peptide LJ-Contacts (L1=purple, MART1=magenta, GVA=orange) and stacked. Error represents SEM over three SMD simulations. Mutants were statisically compared (n=3): #p<0.10, *p<0.05, **p<0.01, ***p<0.001 by one-way ANOVA followed by Tukey-HSD post-hoc test.. Mutants at >0% existence occupancy are statisically compared (n=3) with a Student’s t-test and displayed in red.

### TCR-pMHC Dissociation and Essential Atomic Motion

To assess the mechanism for how the dissociation of the TCR from the pMHC can give rise to a transient strengthening for immunogenic complexes, we investigated essential atomic motion. During the equilibrated state and under load, root mean square fluctuations do not exceed 0.10 nm for any TCR-pMHC interfacial substructure (**Figure 5**). Although there are statistically significant differences in substructure fluctuations between equilibration and pull simulations (i.e., CDR3α, CDR1 β, CDR3β, MHCα, MHCβ, and Peptide), there is no trend except for the peptide. This slight increase for the peptide is expected because as the TCR is dissociated from the pMHC, there is more space for the peptide to fluctuate. Moreover, 0.2 Å differences in fluctuation are unlikely to have physical significance. Additionally, secondary structure analysis of the TCR-pMHC interface under load reveals that all complexes maintain secondary substructures through the bond dissociation process (**Figure S7 Ai-iii, Bi-iii, Ci-iii**). Moreover, a principal component analysis on the TCR-pMHC backbone indicates that more than 96% of essential atomic motion is in the direction of the applied force (**Figure S8 Ai, Bi-iii, Ci-iii, Supplementary Videos**) except for the L1 95 ns (**Figure S8 Aii**) and L1 90 ns (**Figure S8 Aiii**) configurations. Principal component analysis on atomic motion indicates that for these L1 configurations, the TCR-pMHC interface is more energetically stabilized under load than the quaternary structure of the TCR or pMHC, respectively (**Figure S8 Aii-iii, Supplementary Videos)**. This energetic stability results in longer bond lifetime (**Figure S9**) and more transient interactions (**Figure 3–4, Figure S5**). Comparable interfacial fluctuations (**Figure 5**) demonstrate that differences in bond strength are mostly attributed to differences in the equilibrated structures. This is supported by the evidence that the peptide alters the structure of the MHC α-helices, which determines TCR-pMHC bond strength (**Figure 3**).

**Figure 5:**
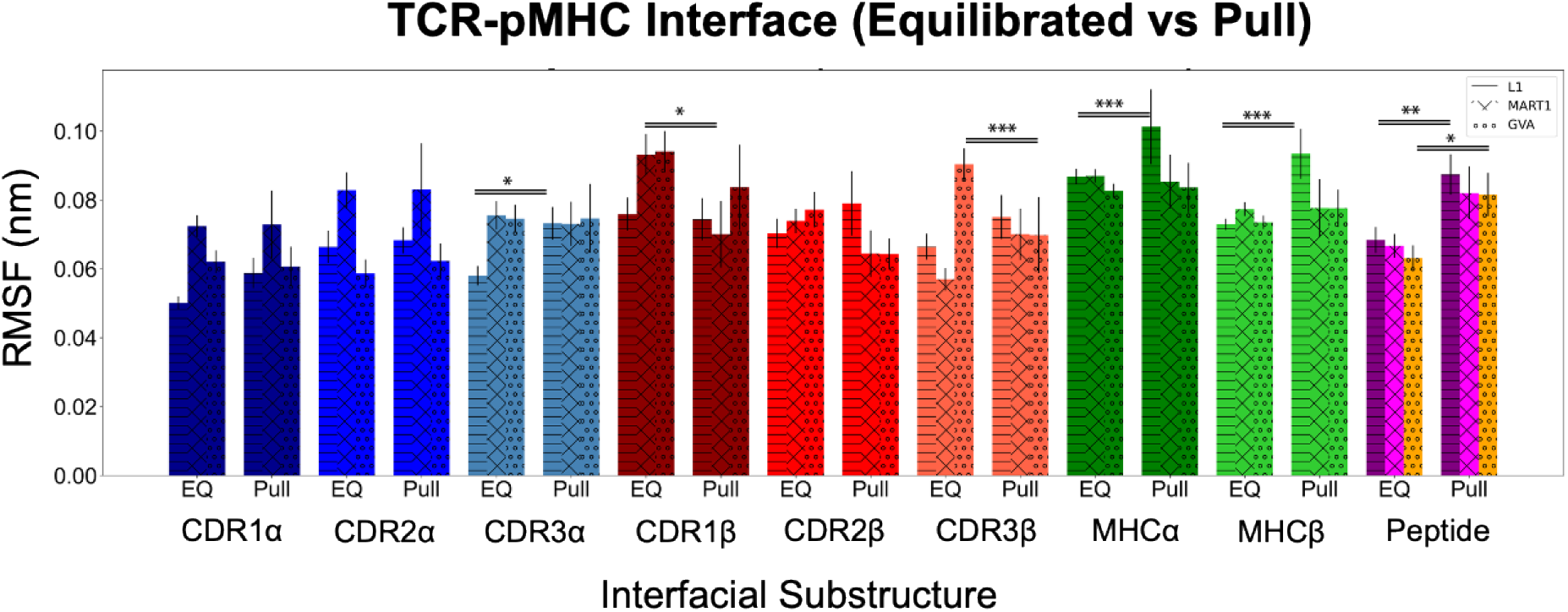
Equilibrated (EQ) and SMD (Pull) interfacial substructure root mean square fluctuations (RMSF) for each TCR-pMHC. EQ error represents SEM on substructure atoms from 90 to 100 ns. SMD error represents SEM on substructure atoms from 3 independent SMD simulations (100, 95, and 90 ns starting configurations). Equilibration and pull subsctructures of mutants were statisically compared: #p<0.10, *p<0.05, **p<0.01, ***p<0.001 by one-way ANOVA followed by Tukey-HSD post-hoc test.

### TCR-pMHC Signature Interaction Maps

To more precisely characterize the alterations, existence maps were created for H-Bonds and LJ-Contacts. The hydrogen bonding maps were generated to include all H-Bonds with greater than 5% existence occupancy and located in at least one of the nine SMD simulations. Furthermore, TCR-MHC and TCR-Peptide interactions were separated to provide the ability to independently examine TCR-MHC and TCR-Peptide hydrogen bonding signatures. To evaluate the precise H-Bond existence as a function of distance, the time axis was converted to COM distance by distributing results into 0.5 Å bins and calculating the fractional occupancy in each respective bin. These H-Bond maps as a function of distance were averaged across triplicate equilibration fluctuations revealing unique hydrogen bonding signatures (**Figure 6 A-C**). The identical TCR-MHC y-axis (**Figure S9A-B**) allows comparison between the three peptides and provides insight into specific hydrogen bonds that may be of particular importance in T cell activation (**Fig. 4A**). For example, the two donated hydrogen bonds donated from TCRα Arginine 27 are absent for GVA, present through reaction coordinate ~ 7.50 nm for MART1, and present through reaction coordinate ~8.30 nm and at greater occupancy for L1 (**Fig. 6**). The maps for each of the three independent simulations for each peptide is provided in the supplement (**Figure S10**).

**Figure 6:**
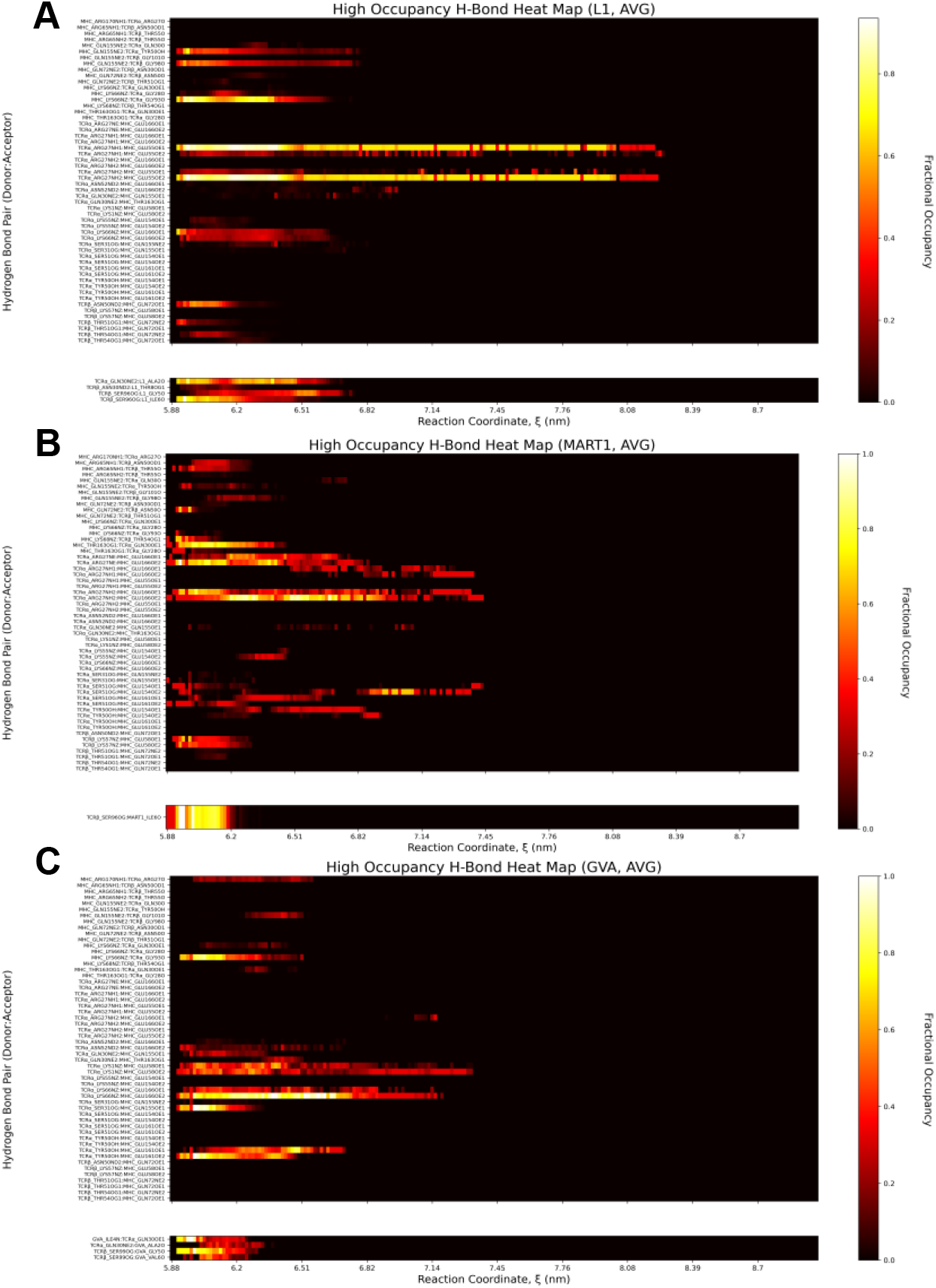
Average hydrogen bond existence maps as function of TCR-pMHC COM distance. The time axis is converted to COM distance by distributing time points into ~0.5 Å bins and calculating the fractional occupancy in each respective bin. The 100, 95, and 90 ns configuration maps of each peptide are arithmetically averaged. The fractional occupancy is represented by the heat scale on the y-axis (right). Peptides **(A)** L1, **(B)** MART1, and **(C)** GVA are comprised of three SMD simulations. The hydrogen bond acceptor and donor are specified on the y-axis (left). Hydrogen bonds are split into interactions between the TCR-MHC (top) and TCR-Peptide (bottom). For TCR-MHC interactions, donor-acceptor pairs with greater than 5% existence occupancy in at least 1/9 simulations are included. For TCR-Peptide interactions, donor-acceptor pairs with greater than 5% existence occupancy in at least 1/3 simulations (for each respective peptide) are included.

Similarly, LJ-Contact maps were generated to include LJ-Contacts with greater than 80% existence occupancy and located in at least one of the nine SMD simulations. Likewise, there is consistency across equilibrated fluctuations and unique sets of LJ-Contacts for L1, MART1, and GVA (**Figure S11**). Moreover, the importance of LJ-Contacts for the L1 peptide (**Figure 4D**) is demonstrated by the nine contacts with over 80% existence occupancy compared to zero for MART1 and GVA (**Figure S11A-I**).

Finally, intra-mutant bond lifetime heterogeneity may depend on the existence of interactions from the starting configuration. For L1, the donated hydrogen bond of TCRα Lysine 66 is present in simulations with ~40 ns bond lifetime and absent in the simulation ~4 ns bond lifetime (**Figure S10A-C)**. Likewise, for MART1, the donated hydrogen bond of TCRβ Lysine 57 is present in the simulations with ~4 ns bond lifetime and absent in the simulation with ~1.5 ns bond lifetime (**Figure S10D-F)**. These results underscore the importance of performing independent simulations with different starting configurations. Additionally, to evaluate the precise bond/contact existence as a function of distance, the time axis is converted to COM distance by distributing results into 0.5 Å bins and calculating the fractional occupancy in each respective bin (**Figure S12A-I, S13A-I**). These maps provide detailed spatiotemporal information on the subsets of interactions that occur at characteristic times and distances.

## CONCLUSION

Our results demonstrate that TCR-pMHC bond strength corresponds with immunogenicity. Bond strength is characterized by stabilized bonds/contacts that mediate formation of transient bonds/contacts at longer times and reaction distances. Interestingly, these bonds/contacts are determined by peptide-mediated conformational shifts in the MHC binding groove. These conformational changes result in unique TCR-MHC interaction signatures. The localized energetics indicate that H-Bonds are crucial for TCR-MHC interactions whereas LJ-Contacts are more consequential for TCR-Peptide interactions. Furthermore, both H-Bonds and LJ-Contacts contribute to overall bond strength and their collective synergy corresponds with immunogenic response. An understanding of fundamental TCR design principles may inform computational strategies to enhance TCR immunogenicity towards novel peptide targets. For example, identifying physiochemical features of the TCR-pMHC interaction that correlate with immunogenicity or targeting residues from the interaction maps may ifnorm TCR mutagenesis strategies. Moreover, this may be a potential approach to overcome the daunting experimental task of matching TCRs to pMHC targets from sequenced repertoires(66, 67). Continued advancements in TCR design principles will provide a discovery platform with broad application in immune-oncology and infectious disease.

## Supporting information

supplemental tables and graphs

## AUTHOR CONTRIBUTIONS

ZAR performed the simulations, analyzed and interpreted the data, and wrote the manuscript. RF designed the experiments, analyzed and interpreted the data, wrote the manuscript, and secured computer time. SCG designed the experiments, analyzed and interpreted the data, wrote the manuscript, and secured the funding.

## ACKNOWLEDGEMENTS

Simulations were performed on the hpc1/hpc2 clusters in the UC Davis, College of Engineering. This work was supported in part by startup funding to SCG from the Department of Biomedical Engineering.

## DATA AVAILABILITY

Starting configurations and Supplementary Videos have been made available: https://doi.org/10.25338/B8FK8D. Custom scripts relevant to the production of figures have been made available on a Github repository: https://github.com/zrollins/TCR-pMHC.git.

