## supplemental tables and graphs for "T Cell Receptor Non-Equilibrium Kinetics"

**SUPPLEMENTARY INFORMATION**

**Figure S1.** TCR-pMHC artificial unfolding due to high force pull from termini

**Figure S2.** TCR-pMHC substructure root mean square fluctuations of 500 pN and 10 pN pulls along the reaction coordinate

**Figure S3.** TCR-pMHC equilibration and root mean square deviations

**Figure S4.** Equilibrated and aligned TCR structures

**Figure S5.** Total H-Bonds and LJ-Contacts between TCR-MHC and TCR-Peptide

**Figure S6.** TCR-pMHC simulated bond lifetime

**Figure S7.** SMD simulations designated secondary structure of TCR-pMHC interfacial substructures

**Figure S8.** Principal component analysis on SMD trajectories

**Figure S9.** List of SMD intersecting TCR-MHC interactions

**Figure S10-S13.** Interaction maps for SMD simulations.

**PC1-PC5.** The starting configurations and trajectories for SMD simulations as well as the projected principal component trajectories have been rendered into videos available at https://doi.org/10.25338/B8FK8D.

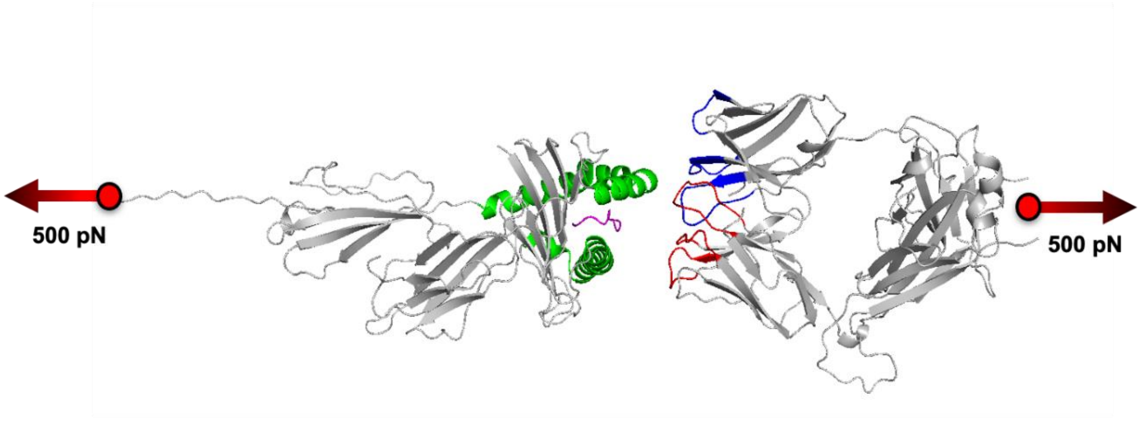


***Figure S5:*** *500 pN pull simulation of TCR-pMHC by terminal residues demonstrating artificial unfolding of pMHC. Linear potential was applied (indicated by red arrows) to the COM of residue 275 of MHC$\alpha$ (left) and the COM of residues 199 and 242 of TCR$\alpha$ and TCR$\beta$, respectively (right)*

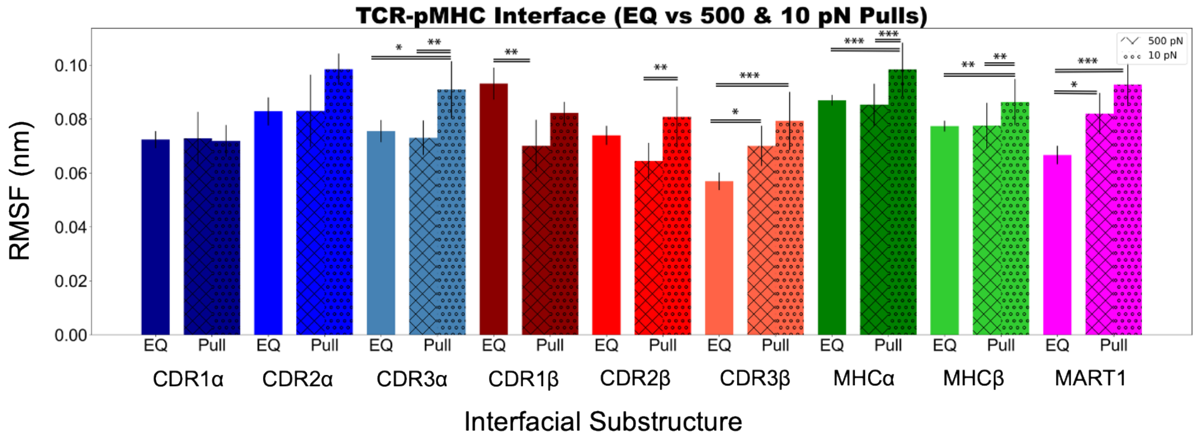


***Figure S2:*** *Interfacial substructure root mean square fluctuations of equilibration (EQ), 500 pN pull, and 10 pN pull. EQ error represents SEM on substructure atoms from 90 to 100 ns. 500 pN pull error represents SEM on substructure atoms from 3 independent SMD simulations (100, 95, and 90 ns starting configurations). 10 pN pull error represents SEM on substructure atoms from 2 independent 50 ns SMD simulations. Equilibration and pull (500 & 10 pN) substructures of MART1 were statistically compared: #p<0.10, *p<0.05, **p<0.01, ***p<0.001 by one-way ANOVA followed by Tukey-HSD post-hoc test.*

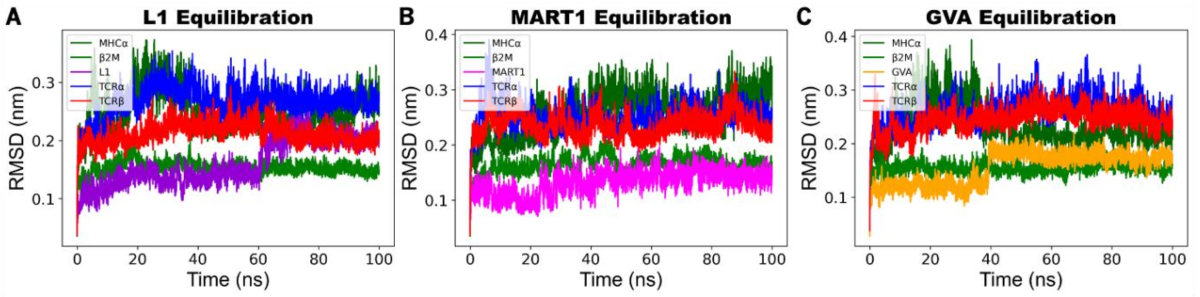


***Figure S3:*** *Equilibration of the TCR-pMHC systems is represented by calculation of root mean square deviation (RMSD) from the initial configuration. Each system is partitioned into 5 protein chains where the TCR is composed of the $\alpha$ & $\beta$ chains and the pMHC is composed of the MHC$\alpha$ chain, $\beta$-2-Microglobulin, and peptide (L1 = purple, MART1 = magenta, GVA = orange).*

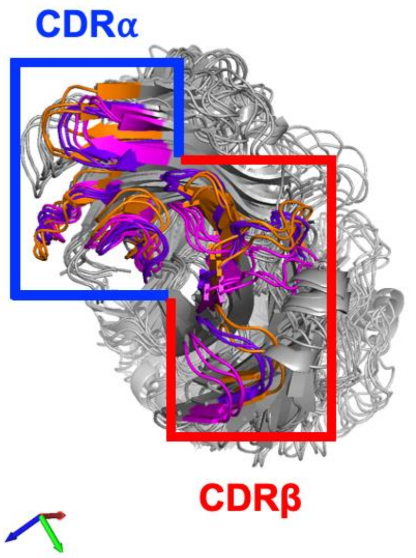


***Figure S4:*** *The equilibrated TCR structures are aligned for each peptide at 100, 95, and 90 ns (L1=purple, MART1=magenta, GVA=orange). The equilibrated CDR$\alpha$ & CDR$\beta$ are indicated in the peptide color to distinguish structure (previously blue & red, respectively, in Fig. 2A-C)*

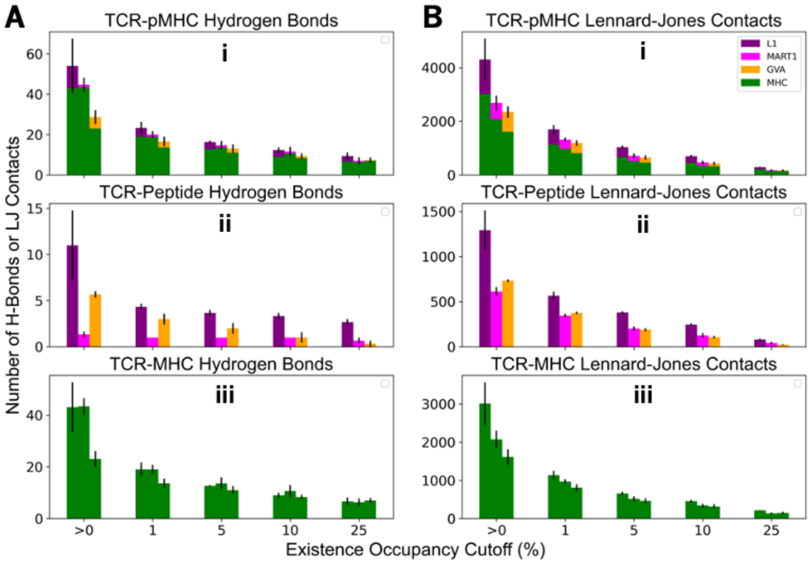


***Figure S5:*** *Hydrogen Bonds and Lennard-Jones Contacts. (A) Number of unique hydrogen bonds are graphed with increasing existence occupancy between (i) TCR-pMHC, (ii) TCR-peptide, and (iii)TCR-MHC. Error represents SEM over three SMD simulations. (B) Number of unique LJ Contacts are graphed with increasing existence occupancy between (i) TCR-pMHC, (ii) TCR-peptide, and (iii) TCR-MHC. Error represents SEM over three SMD simulations.*

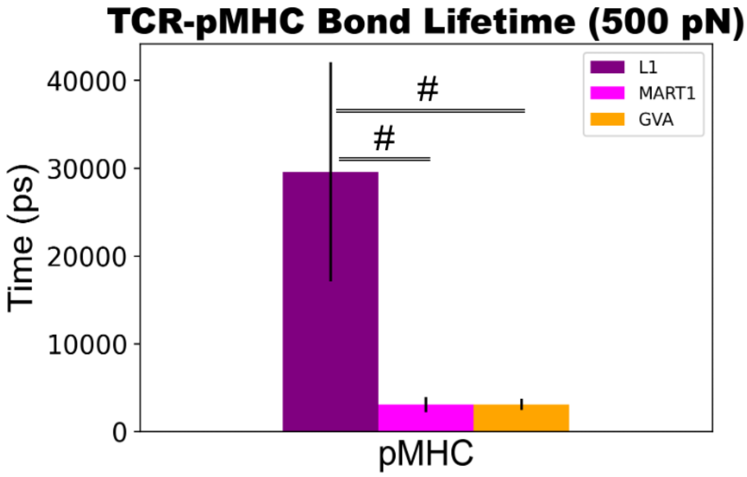


***Figure S6:*** *TCR-pMHC Bond Lifetime. Colors are representative of peptide (L1 = purple, MART1 = magenta, and GVA = orange). Error represents SEM over three SMD simulations. Mutants were statistically compared: #p<0.10, *p<0.05, **p<0.01, ***p<0.001 by one-way ANOVA followed by Tukey-HSD post-hoc test.*

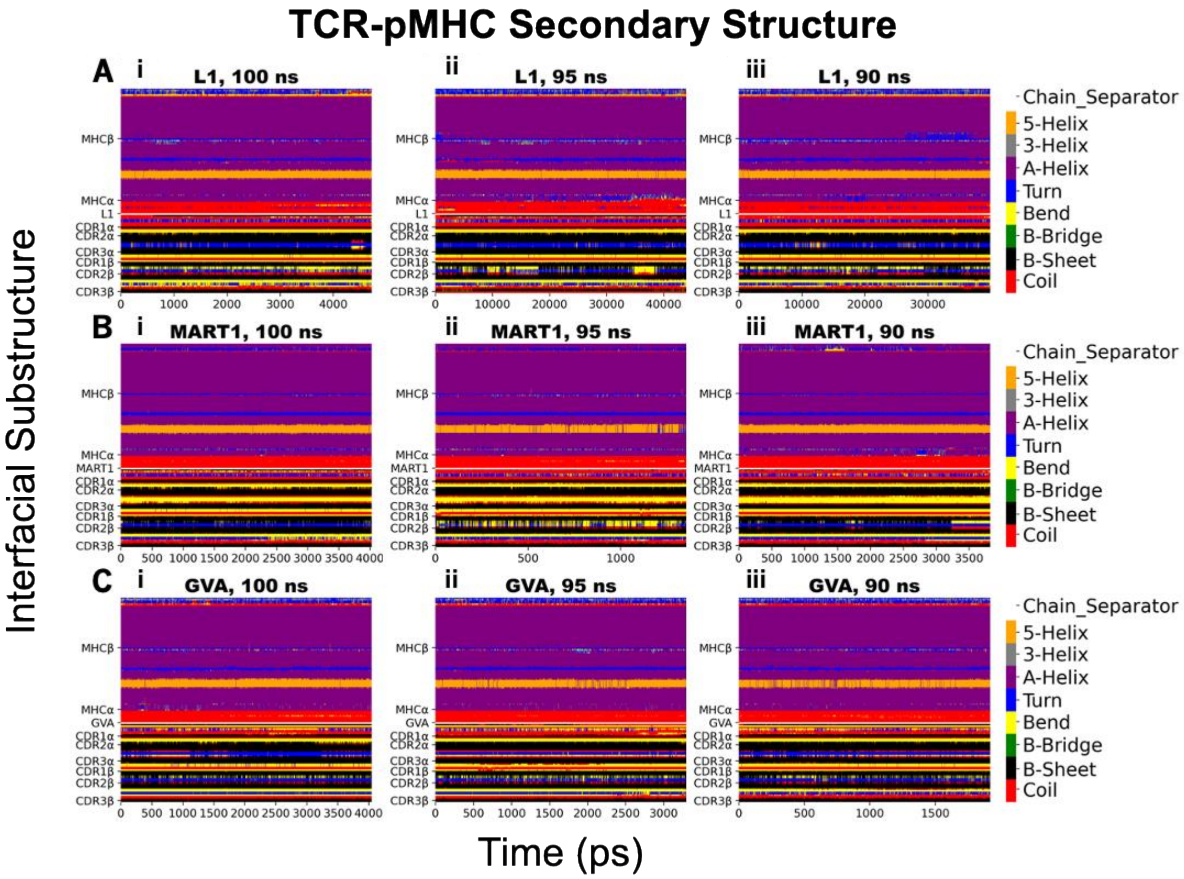


***Figure S7:*** *Designated secondary structure of the TCR-pMHC interfacial substructures for all SMD simulations. Each substructure contains a set of residues (see Methods) located above the demarcation and each residue is assigned a secondary structure over the duration of the simulation. Peptides (A) L1, (B) MART1, and (C) GVA are comprised of three SMD simulations based on their starting configuration: (i) 100 ns, (ii) 95 ns, and (iii) 90 ns.*

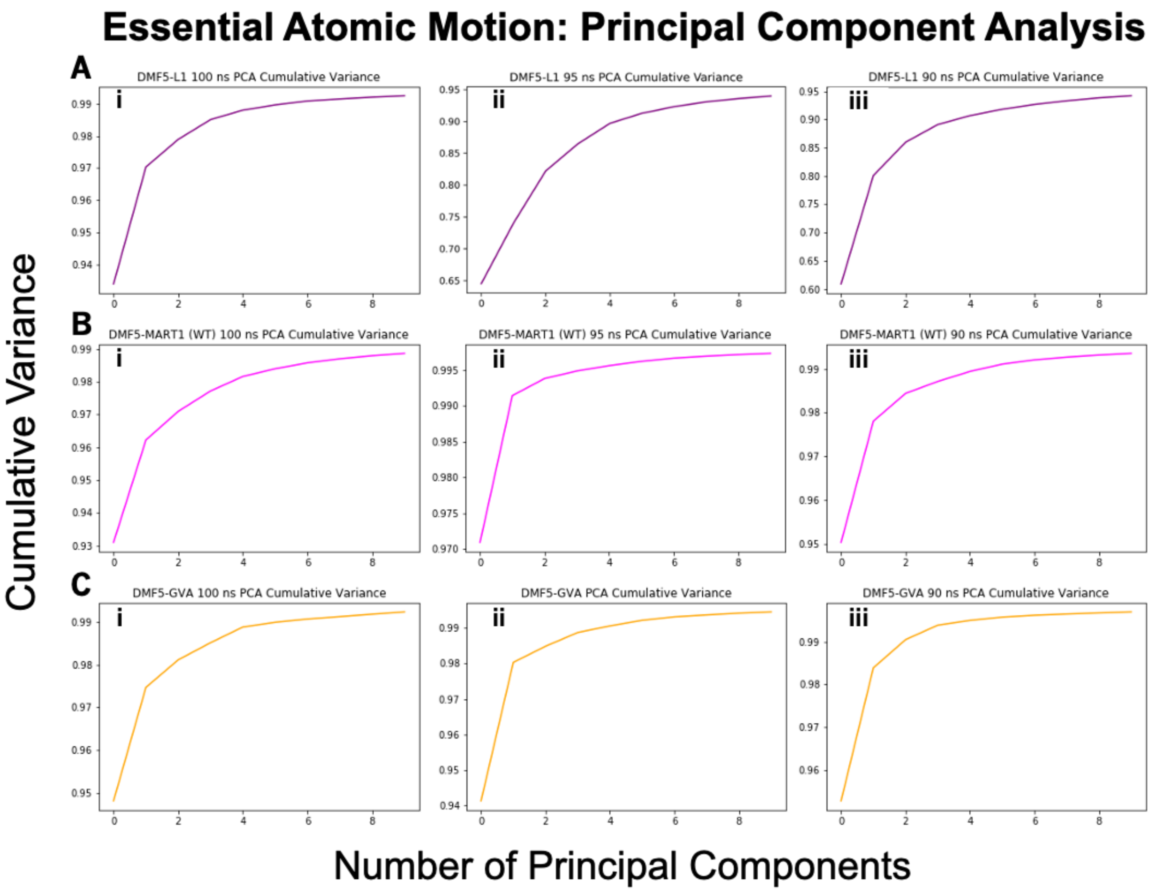


***Figure S8:*** *Principal component analysis was performed on the TCR-pMHC backbone for all SMD simulations. Peptides (A) L1, (B) MART1, and (C) GVA are comprised of three SMD simulations based on their starting configuration: (i) 100 ns, (ii) 95 ns, and (iii) 90 ns.*

**A**

MHC_ARG170NH1:TCRα_ARG27O
MHC_ARG65NH1:TCRβ_ASN50OD1
MHC_ARG65NH1:TCRβ_THR55O
MHC_ARG65NH2:TCRβ_THR55O
MHC_GLN155NE2:TCRα_GLN30O
MHC_GLN155NE2:TCRα_TYR50OH
MHC_GLN155NE2:TCRβ_GLY101O
MHC_GLN155NE2:TCRβ_GLY98O
MHC_GLN72NE2:TCRβ_ASN30OD1
MHC_GLN72NE2:TCRβ_ASN50O
MHC_GLN72NE2:TCRβ_THR51OG1
MHC_LYS66NZ:TCRα_GLN30OE1
MHC_LYS66NZ:TCRα_GLY28O
MHC_LYS66NZ:TCRα_GLY93O
MHC_LYS68NZ:TCRβ_THR54OG1
MHC_THR163OG1:TCRα_GLN30OE1
MHC_THR163OG1:TCRα_GLY28O
TCRα_ARG27NE:MHC_GLU166OE1
TCRα_ARG27NE:MHC_GLU166OE2
TCRα_ARG27NH1:MHC_GLU166OE1
TCRα_ARG27NH1:MHC_GLU166OE2
TCRα_ARG27NH1:MHC_GLU55OE1
TCRα_ARG27NH1:MHC_GLU55OE2
TCRα_ARG27NH2:MHC_GLU166OE1
TCRα_ARG27NH2:MHC_GLU166OE2
TCRα_ARG27NH2:MHC_GLU55OE1
TCRα_ARG27NH2:MHC_GLU55OE2
TCRα_ASN52ND2:MHC_GLU166OE1
TCRα_ASN52ND2:MHC_GLU166OE2
TCRα_GLN30NE2:MHC_GLN155OE1
TCRα_GLN30NE2:MHC_THR163OG1
TCRα_LYS1NZ:MHC_GLU58OE1
TCRα_LYS1NZ:MHC_GLU58OE2
TCRα_LYS55NZ:MHC_GLU154OE1
TCRα_LYS55NZ:MHC_GLU154OE2
TCRα_LYS66NZ:MHC_GLU166OE1
TCRα_LYS66NZ:MHC_GLU166OE2
TCRα_SER31OG:MHC_GLN155NE2
TCRα_SER31OG:MHC_GLN155OE1
TCRα_SER51OG:MHC_GLU154OE1
TCRα_SER51OG:MHC_GLU154OE2
TCRα_SER51OG:MHC_GLU161OE1
TCRα_SER51OG:MHC_GLU161OE2
TCRα_TYR50OH:MHC_GLU154OE1
TCRα_TYR50OH:MHC_GLU154OE2
TCRα_TYR50OH:MHC_GLU161OE1
TCRα_TYR50OH:MHC_GLU161OE2
TCRβ_ASN50ND2:MHC_GLN72OE1
TCRβ_LYS57NZ:MHC_GLU58OE1
TCRβ_LYS57NZ:MHC_GLU58OE2
TCRβ_THR51OG1:MHC_GLN72NE2
TCRβ_THR51OG1:MHC_GLN72OE1
TCRβ_THR54OG1:MHC_GLN72NE2
TCRβ_THR54OG1:MHC_GLN72OE1

**B**

MHC_ALA158HA:TCRα_TYR50CD2
MHC_ALA158HA:TCRα_TYR50CE2
MHC_ALA69O:TCRβ_PHE97HE1
MHC_ARG169HE:TCRα_ARG27HH21
MHC_ARG169HH21:TCRα_ARG27HE
MHC_ARG169HH21:TCRα_ARG27HH21
MHC_GLN155HE21:TCRα_TYR50CZ
MHC_GLN155HE21:TCRα_TYR50OH
MHC_GLN155HE22:TCRα_SER31HG1
MHC_GLN155HE22:TCRβ_GLY98O
MHC_GLN155HE22:TCRβ_THR99HA
MHC_GLU161CD:TCRα_TYR50HH
MHC_GLU166CD:TCRα_ARG27HE
MHC_GLU166CD:TCRα_ARG27HH21
MHC_GLU166OE1:TCRα_ARG27HE
MHC_GLU166OE1:TCRα_ARG27HH21
MHC_GLU166OE1:TCRα_ARG27NH2
MHC_GLU166OE1:TCRα_LYS66NZ
MHC_GLU166OE2:TCRα_ARG27HE
MHC_GLU166OE2:TCRα_ARG27HH21
MHC_GLU166OE2:TCRα_ARG27NE
MHC_GLU166OE2:TCRα_LYS66HZ3
MHC_GLU166OE2:TCRα_LYS66NZ
MHC_GLU55CD:TCRα_ARG27HH12
MHC_GLU55CD:TCRα_ARG27HH22
MHC_GLU55CD:TCRα_ARG27NH1
MHC_GLU55OE1:TCRα_ARG27HH11
MHC_GLU55OE1:TCRα_ARG27HH12
MHC_GLU55OE1:TCRα_ARG27HH22
MHC_GLU55OE1:TCRα_ARG27NH1
MHC_GLU55OE2:TCRα_ARG27HH12
MHC_GLU55OE2:TCRα_ARG27HH22
MHC_GLU55OE2:TCRα_ARG27NH2
MHC_GLY162HA2:TCRα_TYR50HD2
MHC_LYS66HA:TCRα_GLY94HA1
MHC_LYS66HE1:TCRα_GLY28O
MHC_LYS66HG1:TCRα_GLY94HA1
MHC_TYR59HB1:TCRα_ARG27HH12
MHC_TYR59HB1:TCRα_ARG27NH1
MHC_TYR59HD1:TCRα_ARG27HB1
MHC_TYR59HD1:TCRα_ARG27HD2
MHC_TYR59HE1:TCRα_ARG27HB2

***Figure S9:*** *List of TCR-pMHC simulation intersecting interactions (donor:acceptor) for (A) H-Bonds with greater than 5% existence occupancy and (B) LJ-Contacts with greater than 80% existence occupancy*

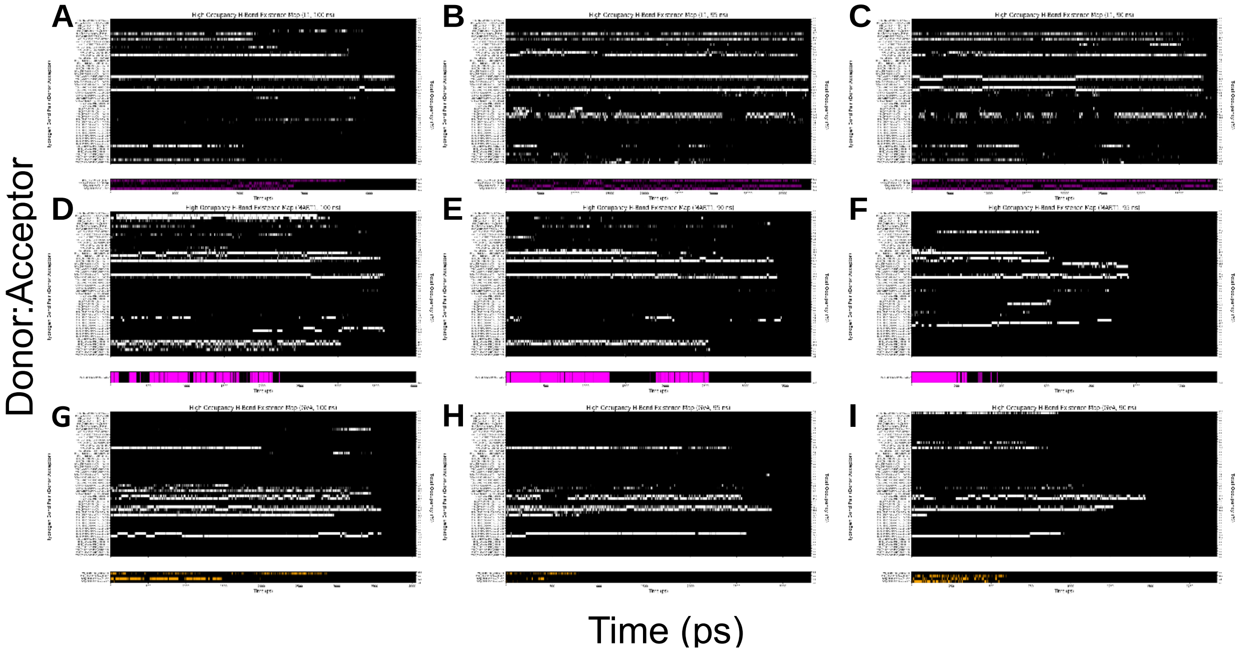


***Figure S10:*** *Hydrogen bond existence maps as function of simulation time. Peptides (A-C) L1, (D-F) MART1, and (G-I) GVA are comprised of three SMD simulations based on their starting configuration: 100 ns, 95 ns, and 90 ns. The hydrogen bond acceptor and donor are specified on the y-axis (left) and percent existence occupancy (right). Hydrogen bonds are split into interactions between the TCR-MHC (top) and TCR-Peptide (bottom). For TCR-MHC interactions, donor-acceptor pairs with greater than 5% existence occupancy in at least 1/9 simulations are included. For TCR-Peptide interactions, donor-acceptor pairs with greater than 5% existence occupancy in at least 1/3 simulations (for each respective peptide) are included.*

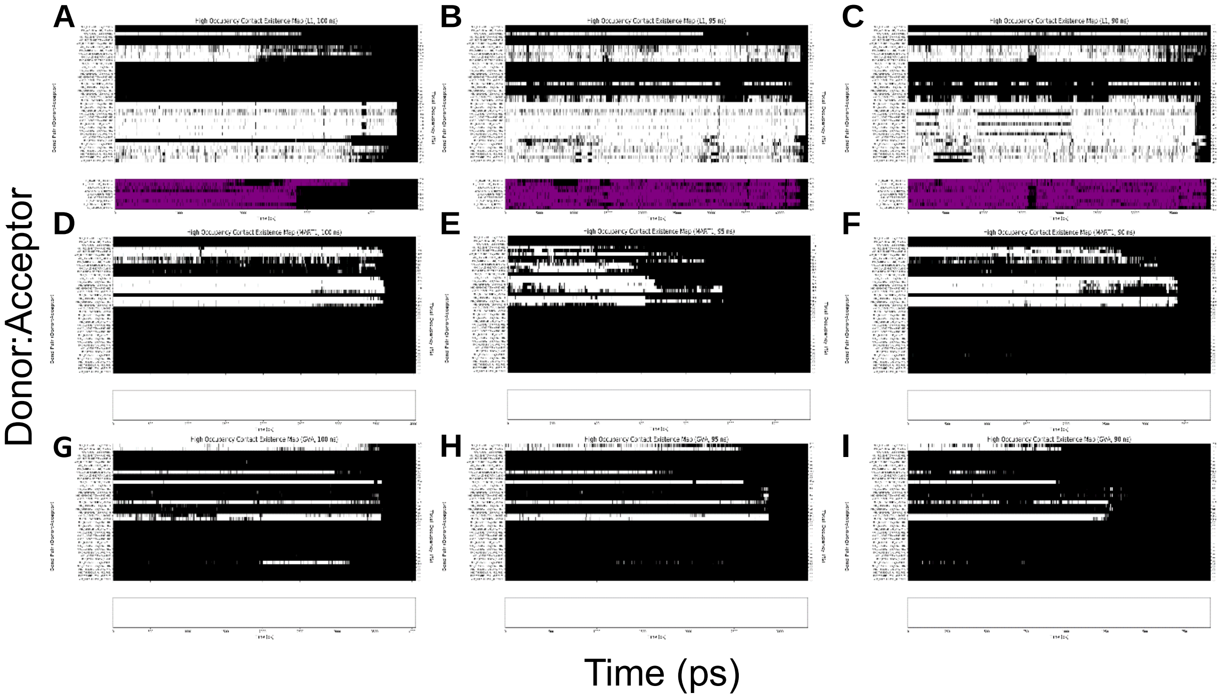


***Figure S11:*** *Lennard-Jones contact existence maps as function of simulation time. Peptides (A-C) L1, (D-F) MART1, and (G-I) GVA are comprised of three SMD simulations based on their starting configuration: 100 ns, 95 ns, and 90 ns. The LJ-Contact acceptor and donor are specified on the y-axis (left) and percent existence occupancy (right). LJ-Contacts are split into interactions between the TCR-MHC (top) and TCR-Peptide (bottom). For TCR-MHC interactions, donor-acceptor pairs with greater than 80% existence occupancy in at least 1/9 simulations are included. For TCR-Peptide interactions, donor-acceptor pairs with greater than 80% existence occupancy in at least 1/3 simulations (for each respective peptide) are included.*

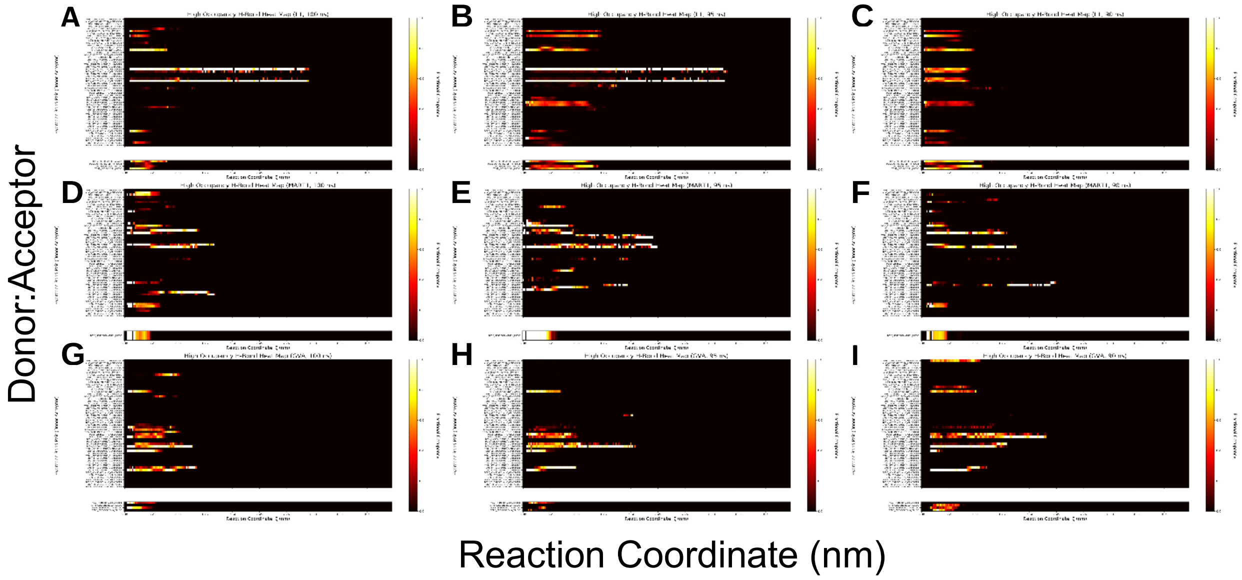


***Figure S12:*** *Hydrogen bond existence maps as function of TCR-pMHC COM distance. The time axis is converted to COM distance by distributing time points into ~0.5 Å bins and calculating the fractional occupancy in each respective bin. This fractional occupancy is represented by the heat scale on the y-axis (right). Peptides (A-C) L1, (D-F) MART1, and (G-I) GVA are comprised of three SMD simulations based on their starting configuration: 100 ns, 95 ns, and 90 ns. The hydrogen bond acceptor and donor are specified on the y-axis (left). Hydrogen bonds are split into interactions between the TCR-MHC (top) and TCR-Peptide (bottom). For TCR-MHC interactions, donor-acceptor pairs with greater than 5% existence occupancy in at least 1/9 simulations are included. For TCR-Peptide interactions, donor-acceptor pairs with greater than 5% existence occupancy in at least 1/3 simulations (for each respective peptide) are included.*

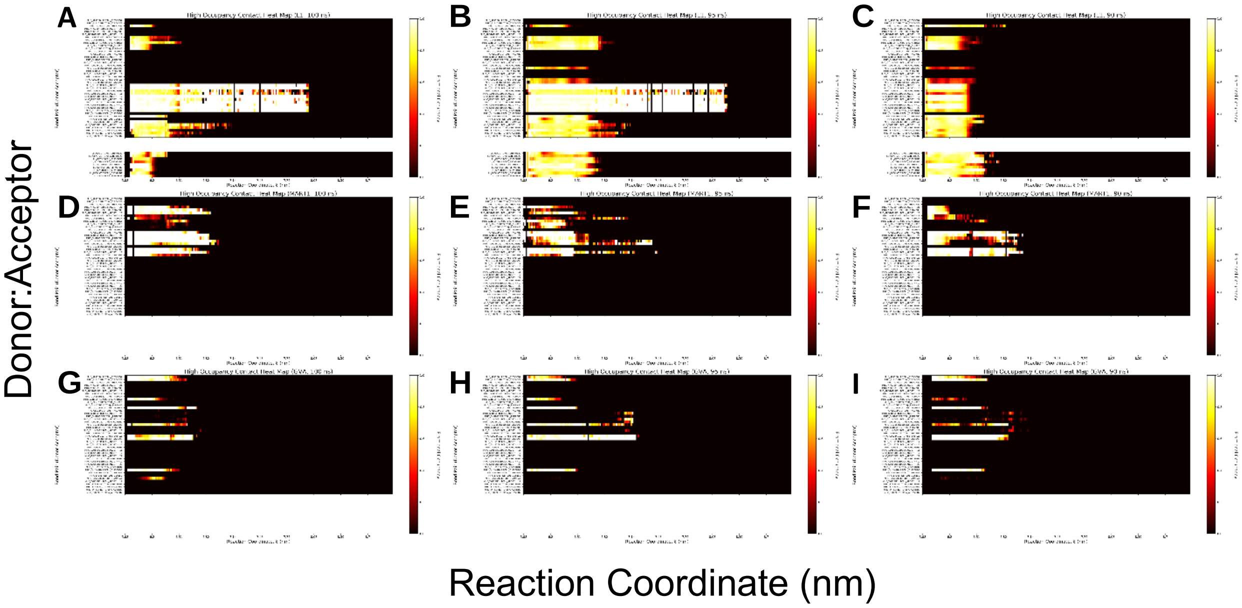


***Figure S13:*** *Lennard-Jones contact existence maps as function of TCR-pMHC COM distance. The time axis is converted to COM distance by distributing time points into ~0.5 Å bins and calculating the fractional occupancy in each respective bin. This fractional occupancy is represented by the heat scale on the y-axis (right). Peptides (A-C) L1, (D-F) MART1, and (G-I) GVA are comprised of three SMD simulations based on their starting configuration: 100 ns, 95 ns, and 90 ns. The LJ-Contact acceptor and donor are specified on the y-axis (left). LJ-Contacts are split into interactions between the TCR-MHC (top) and TCR-Peptide (bottom). For TCR-MHC interactions, donor-acceptor pairs with greater than 80% existence occupancy in at least 1/9 simulations are included. For TCR-Peptide interactions, donor-acceptor pairs with greater than 80% existence occupancy in at least 1/3 simulations (for each respective peptide) are included.*
